# Sparse long-range connections in visual cortex for cost-efficient small-world networks

**DOI:** 10.1101/2020.03.19.998468

**Authors:** Seungdae Baek, Youngjin Park, Se-Bum Paik

## Abstract

The brain performs visual object recognition using much shallower hierarchical stages than artificial deep neural networks employ. However, the mechanism underlying this cost-efficient function is elusive. Here, we show that cortical long-range connectivity(LRC) may enable this parsimonious organization of circuits for balancing cost and performance. Using model network simulations based on data in tree shrews, we found that sparse LRCs, when added to local connections, organize a small-world network that dramatically enhances object recognition of shallow feedforward networks. We found that optimization of the ratio between LRCs and local connections maximizes the small-worldness and task performance of the network, by minimizing the total length of wiring needed for integration of the global information. We also found that the effect of LRCs varies by network size, which explains the existence of species-specific LRCs in mammalian visual cortex of various sizes. Our results demonstrate a biological strategy to achieve cost-efficient brain circuits.

**Highlights:**

- Long-range connections (LRCs) enhance the object recognition of shallow networks
- Sparse LRCs added to dense local connections organize a small-world type network
- Small-worldness of networks modulates the balance between performance and wiring cost
- Distinct LRCs in various species are due to the size-dependent effect of LRCs

**Significance statement:** The hierarchical depth of the visual pathway in the brain is constrained by biological factors, whereas artificial deep neural networks consist of super-deep structures (i.e., as deep as computational power allows). Here, we show that long-range horizontal connections (LRCs) observed in mammalian visual cortex may enable shallow biological networks to perform cognitive tasks that require deeper artificial structures, by implementing cost-efficient organization of circuitry. Using model simulations based on anatomical data, we found that sparse LRCs, when added to dense local circuits, organize “small-world” type networks and that this dramatically enhances image classification performance by integrating both local and global components of visual stimulus. Our findings show a biological strategy of brain circuitry to balance sensory performance and wiring cost in the networks.

**One sentence summary:** Cortical long-range connections organize a small-world type network to achieve cost-efficient functional circuits under biological constraints

## Introduction

Recently, computer vision techniques for object recognition using deep neural networks (DNNs) have been remarkably developed. These now show performance comparable to that of humans^1–5^, implying that deep artificial neural network models can provide collective knowledge of visual processing in the brain^6–8^. However, there exist fundamental differences between the two kinds of systems, particularly regarding the structure required for visual feature abstraction. The brain consists of much shallower hierarchical stages than found in artificial deep neural networks designed for recognizing images with complex feature components. For example, ResNet^3^, one of the state-of-the-art DNNs for object recognition, consists of more than 150 hierarchical layers, thus requires high-level computing power to manage its super-deep structure. However, the ventral visual pathway (from retina to IT) in the brain is composed of less than 10 hierarchical stages^9^ (Fig 1a). This is presumably due to reasons such as the restricted volume of the physical space^10,11^ and limited metabolic resources^12,13^. This raises questions about the distinct strategy implemented in the brain (as compared to artificial networks) that enables cost-efficient visual processing under these constraints.

**Figure.1.**
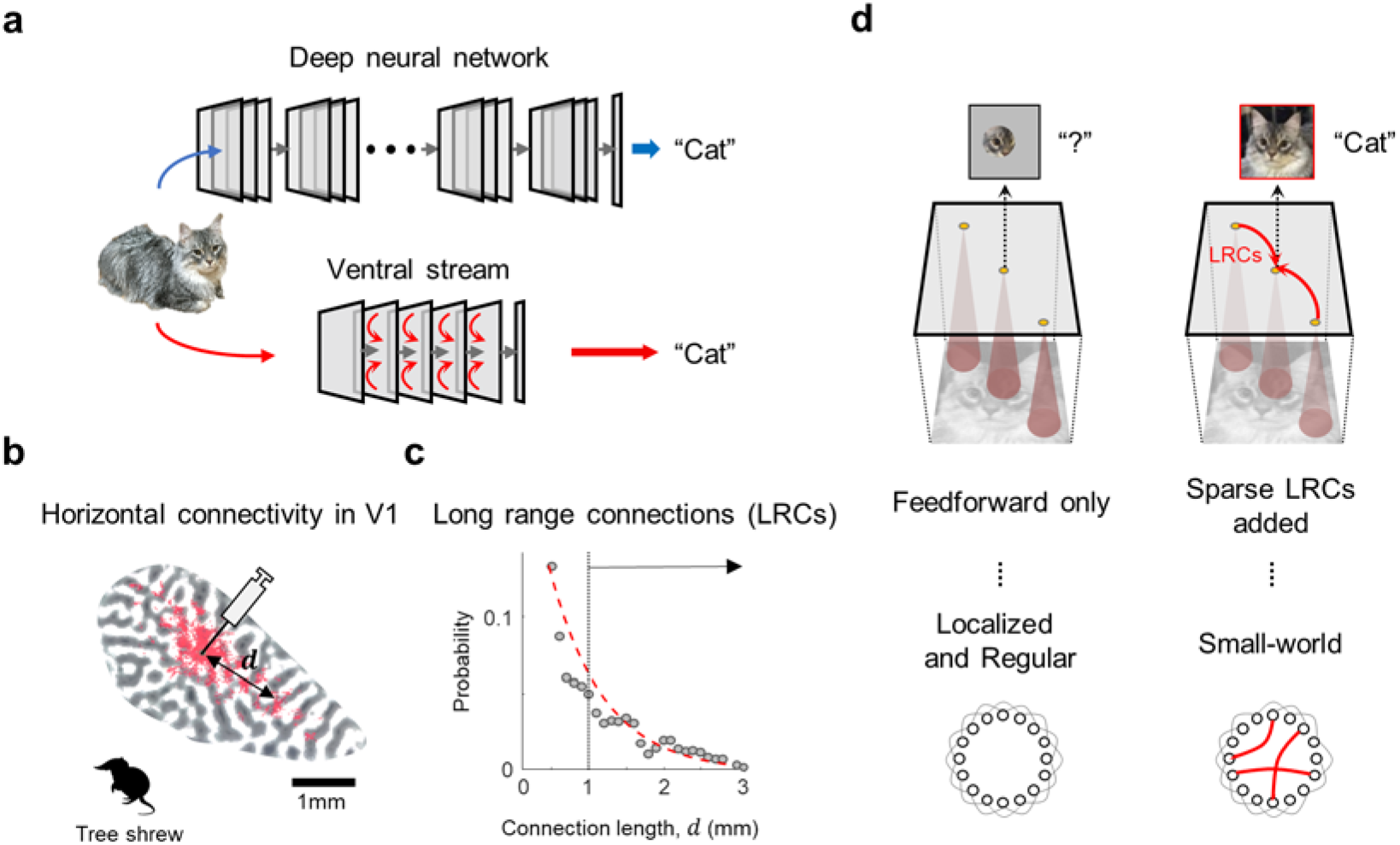
Cortical long-range horizontal connectivity for cost-efficient small-world networks. (a) Hierarchical structures of deep neural networks and the ventral visual pathway. (b) Long-range horizontal connections (LRCs) observed in the visual cortex of tree shrews (redrawn from Bosking 1997, Copyright 1997 Society for Neuroscience). Red dots represent synaptic boutons with retrograde labeling by injection. (c) Distribution of the length of the horizontal connections in tree shrew V1 (gray circle; Bosking 1997). The dashed curve represents an exponential function fitted to data. (d) Illustration of the circuits in shallow networks with and without LRCs. Sparse LRCs added to local feedforward projections organize a “small-world network”.

Long-range horizontal connection (LRC), a distinct anatomical structure observed in the primary visual cortex (V1) of various mammals (Fig 1b) may provide an important insight into this issue. LRCs are found in the visual cortex of monkeys^14^, cats^15,16^, tree shrews^17,18^, gray squirrels^19^, ferrets^20^, and rats^21^. From their extraordinarily long wiring (up to 2-3 mm), LRCs are distinguished from local connections of a short lateral spread (up to 0.5 mm)^22–26^ (Fig 1c). Previous studies have suggested that LRCs may play a particular role in visual processing, compensating for their large wiring cost, and possibly contributing to layer-wide functions such as contextual modulation at early stages^27–29^. However, the exact functions of LRCs for visual processing are still unknown.

Here, we suggest that LRCs added to local cortical connections can organize a “small-world network”^30^ in visual cortex, which enables cost-efficient object recognition under resource constraints (Fig 1d). The small-world, a term first introduced by Watts and Strogatz^30^, is a network of high local clustering and short average global path length. We hypothesized that this may describe a characteristic structure of neural circuits required to encode both local and global feature components of visual images. Particularly in circuits of large V1, we assumed that mere local connections would not be sufficient to organize a small-world network, but LRCs could act as shortcuts for cortical inter-neural communications, organizing the entire V1 circuitry as a “small-world” (Fig 1d). On the other hand, LRCs may not contribute significantly in a small V1 network where local connections suffice.

To validate this idea, we implemented a model neural network of V1 and performed simulations for image classification capability of the network. We found that addition of the LRCs significantly enhances object recognition performance of a shallow network, and that the degree of performance enhancement is strongly correlated to the small-worldness of the circuit. Our key prediction that the effect of LRCs is dependent on the size of the network was validated by our analysis of animal data on the existence of LRCs in diverse mammal species. Taken together, our results provide a comprehensive understanding of the functional role of LRCs in V1.

## Results

### LRCs enhance object recognition performance of shallow networks

To test the contribution of LRCs to object recognition of shallow neural networks, we designed a three-layer convergent neural network as a simplified model of the ventral visual pathway (Fig 2a). The model network consisted of 784 input units, one hidden layer of 784 units, and 10 output units for classification. Neurons in the input and the hidden layers were linked by local feedforward connections, following observed convergent projections of the early visual pathway^31^. Model long-range connections were added in the lateral connections of the hidden layer.

**Figure 2.**
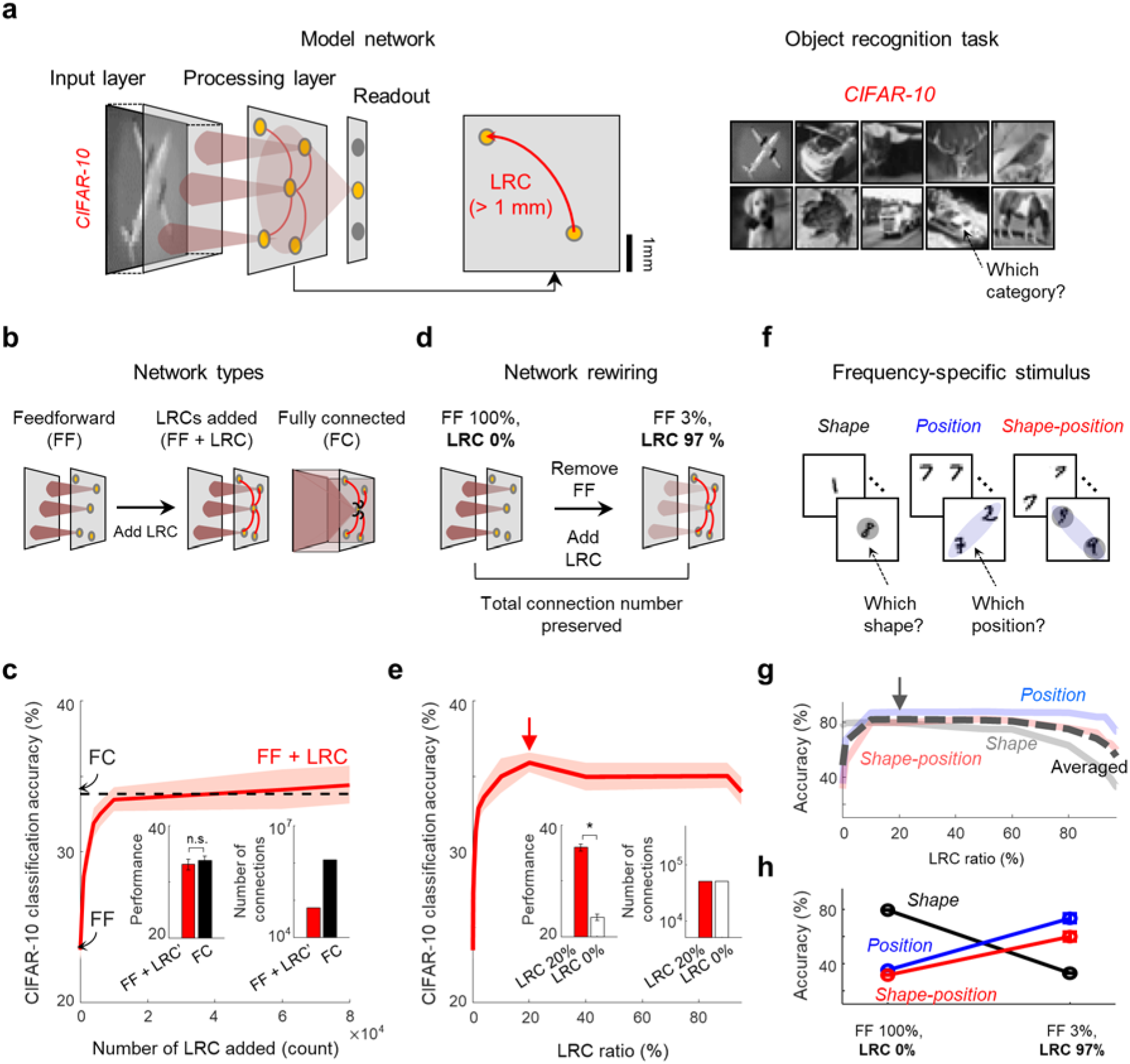
Long-range connections enhance image classification of shallow networks. (a) A simplified model of shallow networks in the visual pathway (left). Examples of the CIFAR-10 dataset used for image classification task (right). (b) Three types of shallow networks with different connectivity. (c) Image classification performance of the networks in (b). LRCs added to local feedforward networks drastically enhance classification accuracy (red solid line), comparable with that of FC (black dashed line). Performance of the FF + LRC (N = 10,000 LRCs) and the FC networks (left inset; Mann-Whitney U-test: p = 0.15). This FF + LRC network uses only 5% of the number of connections in the FC (right inset). (d) The network rewiring process. Feedforward connections (FF) are removed and the same number of LRCs are added to preserve the total number of connections. (e) Classification accuracy for CIFAR-10 datasets during the rewiring process. Red arrow indicates the maximum accuracy achieved. Classification performance increases until the ratio of LRCs reaches 20% of the total number of connections (left inset; Mann-Whitney U-test: *p < 10^−14^), where the amount of total wiring in the network is unchanged (right inset). (f) Modified MNIST datasets for three types of classification test. The “shape” dataset requires identification of local shape of the digits, and the “position” dataset requires classification of the position of digits. The “shape-position” dataset requires both. (g) Classification accuracy for each dataset with variation of the LRC ratio. Average accuracy for all three tasks (black dashed line). Black arrow indicates the maximum accuracy. (h) Classification accuracy for each dataset of the network with 0% LRC and with 97% LRCs. Error bars and shaded area represent the standard deviation of the accuracy for 20 trials.

We trained and tested the model network using CIFAR-10 datasets for natural image classification. Three types of network conditions with different connectivity were compared (Fig 2b): (1) feedforward only (FF), (2) feedforward with LRCs (FF+LRC), and (3) fully-connected (FC). As expected, we found that purely feedforward networks without LRCs show lower classification accuracy (23.5%) (Fig 2c) than that of a fully connected network (33.7%) (Fig 2c, black dashed line) which may indicate the maximum capability of this shallow network. Notably, we found that adding only a small number of LRCs to the FF network (Fig 2c) significantly enhances the network performance, comparable to that of the FC. When the performance of FF+LRC becomes equivalent to that of the FC (Fig 2c, left inset, Mann-Whitney U-test: p = 0.15), it uses only 5% of the number of total learnable connections compared to that of the FC network (Fig 2c, right inset). This indicates that LRCs can noticeably enhance the object recognition of shallow networks in a cost-efficient manner. We also confirmed that the performance improvement by LRCs was fairly consistent across different conditions of feedforward connectivity such as sampling ratio and convergence range (Supplementary Fig 1).

To investigate the cost-efficiency of LRCs more specifically, we varied the ratio of LRCs (the number of LRCs per total number of connections) in the network while the total number of connections were unchanged. We added LRCs while disconnecting the same number of local feedforward connections, and measured changes in the classification performance of the network (Fig 2d). We found that the accuracy was dramatically improved by additional LRCs, and that the performance was maximized when sparse (20%) LRCs were combined with the dense local feedforward network (Fig 2e, brown asterisk) (Fig 2e inset, Mann-Whitney U-test: *p < 10^−14^). This result indicates that a certain portion of LRCs inserted in a shallow FF network can optimize the cost-efficiency of the network.

Next, to investigate the contribution of the LRCs to images of a different spatial frequency spectrum, we generated new datasets of various frequency components (“shape”, “position”, and “shape-position”), by organizing different combinations of MNIST datasets (Fig. 2f, see Supplementary Fig 2 for details). This demands that the network encodes either local features (“shape”), global features (“position”), or both, to classify each class correctly. Using these datasets, we trained the network and examined the classification performance while varying the ratio of the LRCs again. We found that when the LRC ratio was 0%, the network showed higher performance for “shape” datasets than for the other two datasets (Fig. 2g and 2h, LRC ratio = 0%). On the other hand, the network with a large portion of LRCs showed better performance for “position” and “shape-position” datasets (Fig. 2g and 2h, LRC ratio = 97%). In particular, the network with a certain portion (20%) of LRCs showed the maximum average accuracy for the three datasets, implying a consistent classification capability for various types of images (Fig. 2g, brown asterisk). In addition, this result supports our hypothesis that LRCs enhance the network performance by integrating global features of images that cannot be captured by local connections only.

### Combination of LRCs and local connections organizes a small-world type network

It is notable that networks with combined sparse LRCs and dense local connections are advantageous for encoding a wide range of the frequency spectrum of natural images. Interestingly, this connectivity architecture is comparable to a “small-world network^30,32^”, in terms of both structure and purpose. A small-world type network is defined as a network of high local clustering and short average global path length^30^. One is generally achieved using a number of local clusters with sparse long shortcuts (Fig 1d, bottom). Small-world networks have advantages of both high “local” interactions between adjacent nodes and short “global” distance between distant nodes. From this similarity, we hypothesized that a “small-worldness” index would efficiently describe the ability of our network to classify both “local” and “global” components in natural images (Fig 3a).

**Figure 3.**
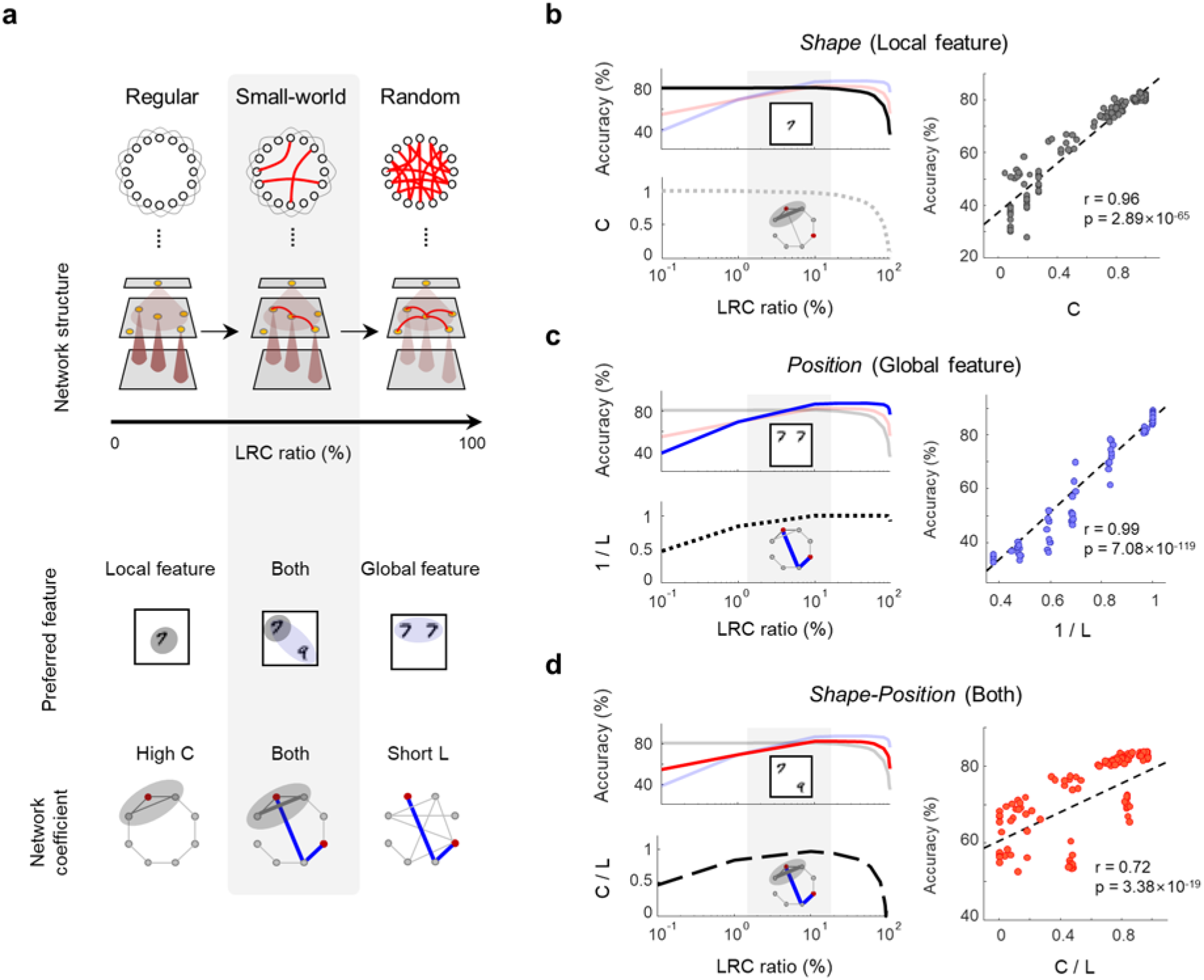
Network-theory approach to long-range connectivity. (a) Addition of LRCs to a local FF network modulates the network connection structure toward a small-world network, enabling correct classification of both local and global features in images. (b) Classification accuracy for “*Shape”* dataset (left top) and the clustering coefficient (C, left bottom) of the network during the rewiring process are strongly correlated (right). (c) Classification accuracy for “*Position”* dataset (left top) and reciprocal of average path length (1/L, left bottom) are also correlated (right). (d) Classification accuracy for “*Shape-Position”* dataset (left top) and the small-worldness index (C / L, left bottom) are correlated significantly (right). Gray shaded area represents “small-world” regime.

To test this hypothesis, we investigated correlations between the classification performance of the model network and a network structure described by the clustering coefficient (C), average path length (L), and small-worldness (SW) index^32^. We found that performance enhancement was strongly correlated to the corresponding network parameters. Change in the clustering coefficient (C) is strongly correlated with performance for a “shape” dataset (Fig 3b), and a reciprocal of the shortest path length (1/L) is strongly correlated with performance for a “position” dataset (Fig 3c). As a result, the small-worldness, a value defined as C divided by L, shows a significant correlation with performance for a “shape-position” dataset (Fig 3d). We confirmed that the addition of LRCs to an FF network increased the image classification performance until the small-worldness reached the maximum. This suggests that the combination of sparse LRCs and dense local feedforward connections allows emergence of a new small-world network, and enables consistent object recognition of various types of images.

### Size-dependency of small-worldness predicts the existence of species-specific LRCs

LRCs are observed in various mammalian species^14–21^, but not in rodents smaller than rats (Fig 4a). Mice and rats are genetically close relatives with neuroanatomical and functional similarities, including organization of functional circuits in V1^33–35^. Despite their genetic proximity, rats have LRCs clearly distinct from their local lateral spread, whereas mice do not (Supplementary Fig 3). Here we propose that the contribution of LRCs to small-worldness varies according to the size of the networks, and that this can explain the existence of species-specific LRCs.

**Figure 4.**
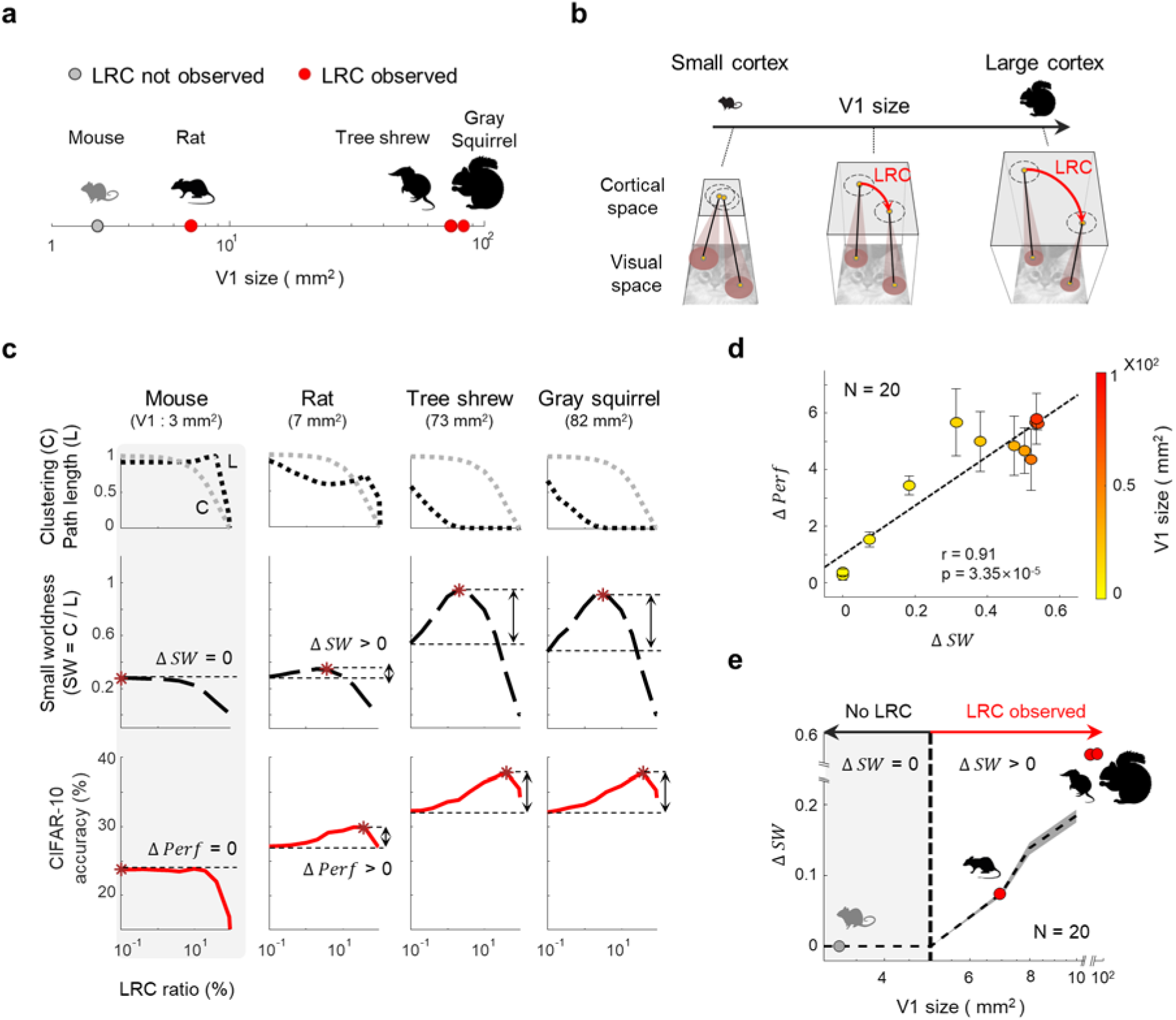
Long-range connections do not increase the small-worldness in small cortices. (a) Species with and without LRCs. (b) Illustration of the size-dependent effect of LRCs. Shaded circles represent local visual area (receptive field) projected onto target cortical neurons indicated as yellow dots. Note that LRCs in a large cortex can link two distant cortical neurons, the receptive fields of which are not overlapped. (c) The CIFAR-10 classification accuracy and the network parameters as a function of LRC ratio (mouse, rat, tree shrew, gray squirrel). Changes in performance accuracy and small-worldness, ΔPerf (accuracy) and ΔSW (small-worldness), were defined as the difference between the maximum and the initial value. Gray dashed lines represent the clustering coefficient (C) and black dashed line indicate the path length (L). Brown asterisks indicate the maximum points of accuracy and small-worldness curves. (d) Correlation between ΔPerf and ΔSW of networks for cortices of various sizes. (e) Changes of small-worldness, ΔSW, as a function of V1 size. Note that ΔSW becomes positive only when the cortex size exceeds 5 mm^2^. Error bars represent the standard deviation for 20 trials in (d) and (e).

Retino-cortical feedforward projections organize a precise retinotopy in most species^34,36,37^, and visual information from a local retinal area is projected onto a local cortical space. If the size of the cortex is large, two distant cortical neurons that receive inputs from distinct retinal spaces cannot communicate because their local connections do not overlap. However, addition of LRCs in this large cortex enables communication of distant neurons by reducing the average path length in cortical space. In this condition, LRCs increase dramatically the small-worldness of a cortical network (as shown in Fig 3b-d). In contrast, if the size of the cortex is relatively small, as in mice, introducing LRCs would not further decrease the path length, because cortical neurons already share information with neighboring neurons via local connections. Thus small-worldness will remain unchanged even though a large number of LRCs are added. This scenario implies that LRCs may not affect the performance of networks of small size.

To validate this prediction, we performed connectivity analysis of networks of various sizes, simultaneously with the image-classification performance test: The CIFAR-10 classification performance was estimated while the small-worldness was measured in model cortical networks of various sizes (Fig 4b). When the model cortex size was large (gray squirrel^35,38^, 82 mm^2^), introduction of LRCs increased both classification accuracy and small-worldness (Fig 4c, rightmost). In contrast, when the cortex size was small (mouse^35,39^, 3 mm^2^), the performance and small-worldness did not increase further regardless of how many LRCs were added (Fig 4c, leftmost). In general, we found a strong correlation between the increment of performance (ΔPerf) and small-worldness (ΔSW) in relation to the variation of V1 size (Fig 4d). Notably, ΔPerf and ΔSW were positive only when the cortex size exceeded a certain threshold (~5 mm^2^, Fig 4e; Supplementary Fig 4). These results provide a simple explanation of the existence of species-specific LRCs.

## Discussion

We showed that cortical long-range connections can play an important role in developing a cost-efficient circuit under structural constraints. In shallow networks with large processing area, LRCs act as shortcuts for the communication of distant neurons, which forms a small-world network. In networks of small area, however, LRCs could not contribute in such ways and this provides a possible explanation for the existence of species-specific LRCs in various mammals.

An ability to recognize natural images is a fundamental function in animals that is required for survival. Because natural images contain a significant portion of low-frequency component^40^, a successful visual circuit should be able to encode this “global” information^41^. This requires sufficiently deep hierarchical structures and a large number of connections^42^, unless the convergent range of feedforward projections is large enough (i.e., comparable to fully connected networks). Any of the scenarios above require a significant amount of wiring cost. However, due to the restricted volume of the physical space and limited metabolic resources, the visual pathway in the brain does not include such a deep structure. This makes it difficult to encode the global features of visual input with localized feedforward projections. In this case, implementing a wide range of convergence between the layers also may not be an appropriate choice because it may lead to a significant loss of high-frequency “local” information. Here, according to our model, the combination of local convergent projections and sparce LRCs may enable the brain to address this issue by capturing both local and global information, using a minimum of wiring cost.

In the current study, for simplicity, we neglected detailed biological parameters except the size of V1 in the analysis of the small-worldness of the network (Fig 4). However, there are a number of other factors that affect the small-worldness condition of networks in the visual pathway: cell density, the convergent range of feedforward projections, the nonlinear magnification factor in the retinotopy, and the recurrent and feedback circuits. These are all possible factors affecting the information processing. Detailed effects of these factors need to be considered in subsequent studies.

An additional question about the formation of LRCs during development remains. It has been suggested that LRCs develop without visual experience, because they are observed before eye-opening^43,44^. This suggests that LRCs, a characteristic architecture that links cortical neurons of similar tuning, may emerge spontaneously without training. Previous studies have reported that feedforward afferents from the periphery may play important roles in early development of the cortex^45,46^, and that LRCs in sensory cortices can be induced by feedforward projections from the retina^47^. In a previous study^48^, which is relevant to the current findings but contains independent findings, we showed that long-range horizontal connections in V1 may emerge from spatio-temporally structured retinal waves generated spontaneously, implying that cortical LRCs may originate from early peripheral activities before eye-opening. Further investigation of this scenario from the analysis of developing V1 in young animals may provide further support for our model.

Overall, our results suggest that cortical long-range connections enable organization of small-world type networks for cost-efficient visual recognition under biological constraints. This finding offers a simple but powerful model that explains the role of cortical long-range connections and an underlying mechanism for species-specific variants. This reveals a biological strategy of the brain to balance functional performance and resource cost.

## Methods

### Neural network model for image classification

All simulations were performed using the MATLAB deep learning toolbox. To implement a simplified model of the ventral visual pathway, we used the multilayer perceptron with three layers, consisting of an input layer of 784 units, one hidden layer of 784 units, and an output layer of 10 units for classification. The schematic architecture of the model is shown in Fig 2a.

The feedforward connectivity between the input and hidden layers were set to local convergent structure, following the study of Hubel and Wiesel^31,49^. The i^th^ neuron in the hidden layer has a circular receptive field, that is

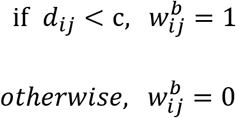

where j is the index of the neuron in the input layer, and *d*_*ij*_ is the Euclidean distance between neurons i and j when the input layer is projected to the hidden layer. Here, c denotes the size of the receptive field, which is equivalent to maximum distance *d*_*ij*_, which was initially set to 4. The term 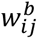 is a Boolean parameter that represents the presence of connection between neuron i and j. Connections between the hidden and output layer were randomly connected with probability 0.5.

Lateral connections could only be added in the hidden layer. The length distribution of lateral connections followed the V1 data^17^ observed in tree shrew V1 (Fig 1c). To match the statistics observed in the biological data, we assumed the condition that the 15° visual images were shown to a head-fixed tree shrew. Considering the magnification factor and V1 size in a tree shrew^17,35,50^, the scaling factor was set to 0.1 mm/unit, so that one unit of the input and hidden layers matched 0.1 mm in the cortical space of the biological data. Among the lateral connections, long-range connections were defined as lateral connections longer than 1 mm^51^.

To perform image classification, our model was trained using stochastic gradient descent, even though it is not strictly a biologically plausible learning rule^52^. The batch size was set to 200 images and the weight decay factor was set to 0. All connection weights were initialized from a zero-mean Gaussian distribution with a standard deviation of 0.01. The learning rate was set to a constant value of 0.1. Other hyperparameters such as the number of epochs were chosen to provide reasonable accuracy in the image classification task.

### Image datasets

The model was trained for performing the image classification task. Depending on the purpose, we used four kinds of image datasets. To test the ability of the model network for overall image recognition performance, the CIFAR-10 dataset^53^ was used as an example of a natural image. Before training the network, the image was converted to greyscale using the MATLAB rgb2gray function.

To investigate the frequency specific role of LRCs for image classification, we generated three datasets by modifying the size and position of handwritten digits (MNIST). Details are as follows:

1. The “shape” dataset was designed by resizing a hand-written digital image to 8 × 8 pixels in the center of 28 × 28 pixels. The “local” dataset consists of eight categories depending on the number (selected from 1 to 8). For this dataset, only local information (shape) of the digits was required for classification.
2. The “position” dataset was made using the following procedure. First, the 28 × 28-pixel area was divided into four 14 × 14 areas. Second, two out of the four areas were chosen. Third, a randomly selected 8 × 8-pixel digit was inserted into a random position in the chosen area. The “global” dataset also consisted of eight categories depending on the position where the digit was located. The type of digit is irrelevant for the classification.
3. The “shape-position” dataset was generated by the following procedure. First, the 28 × 28-pixel area was divided into four 14 × 14 areas. Second, two diagonally aligned areas were chosen (either 45° or 135°). Third, an 8 × 8 pixel digital image (either “7” or “9”) was located in a random position of the selected area. The reason we chose “7” and “9” among ten numbers was to make the difficulty of the task match that of the previous tasks. The “shape-position” dataset consisted of eight categories depending on the type and location of the digits.

For the “shape” dataset, the local shape of the digits was the only information related to the classification. For the “position” dataset, the shape of the numbers is irrelevant and only the position where two numbers are placed was important information for the classification. For the “shape-position” dataset, the network should perceive both the shape and position of the digits in order to classify the images correctly. For all three datasets, each category contained 1000 generated images. We properly tuned the difficulty of the three tasks to be similar by adjusting the background noise level and position of the numbers. This worked because they did not have to be exactly the same.

### Test of image recognition performance according to the connectivity

To examine the exact role of a long-range lateral connection (Fig 2), we varied the ratio of the LRCs in the network and tested the image recognition performance after training of the modified MNIST dataset (shape, position, shape-position). Note that, in order to fairly compare the performance of networks that had different structures, the total number of connections had to be controlled because it is obvious that the network that has more learnable parameters should perform better. To keep the number of learnable parameters the same while varying the network connectivity, we ablated the feedforward connections between the input and hidden layer while adding the same number of LRCs (described in Fig 2b). The connections between the hidden and output classification layer were not ablated because those connections were thought to be required for the classification, but not for the image abstraction.

### Network connectivity indices

Overall, the network parameters we used in the paper followed earlier definitions^32^ with small modifications. The clustering coefficient *C* is the average density of connections between the neighbors and is described as

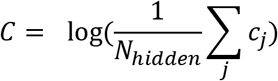

where *N*_*hidden*_ the number of hidden neurons and *c*_*j*_ is the local coefficient of *j* hidden neurons. Here, *c*_*j*_ is determined by the following equation:

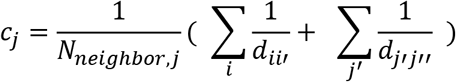

where *N*_*neighbor,j*_ is the number of neurons connected with *j* hidden neurons, *d*_*ii′*_ is the inter-neighbor distance between source *i* input neuron and destination *i*’ input neuron, which are connected with *j* hidden neuron. Similarly, *d*_*j′j*”_ is the inter-neighbor distance between source *j*′ hidden neuron and destination *j*′ hidden neuron, which are connected with *j* hidden neuron.

The characteristic path length *L* si defined as:

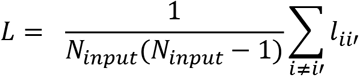

where *N*_*input*_ is the number of input neurons and *l*_*ii*′_ is the shortest path between *i* and *i*′. The term *l*_*ii*′_ is defined as the shortest path from a random input neuron I to another input neuron *i*′. Please note that the shortest path must be include at least one hidden neuron because the network is a layered structure.

The small-worldness was defined as the ratio of the clustering coefficient C and path length L. However, this index is largely influenced by network parameters such as the number of units. To avoid this problem, the small-worldness SW was modified to the ratio of the normalized clustering coefficient C and normalized path length L as follows:

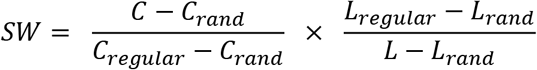

where *C*_*rand*_ and *L*_*rand*_ are the clustering coefficient and characteristic path length on a random network consisting of only randomly generated LRCs. The terms *C*_*regular*_ and *L*_*regular*_ are the clustering coefficient and characteristic path length on a random network consisting of only FFs.

### Network model for various species

To model the visual pathways of four different species (mouse, rat, tree shrew, and gray squirrel), we varied the size of the processing layer (also the number of neurons) according to the relative V1 size of each species. All the other parameters (including input cell density, sparsity of feedforward connections, and V1 cell density) were assumed to be consistent across species. The anatomical data of averaged V1 size used in this study were from mouse^35,39^ (3 mm^2^), rat^35,54^ (7 mm^2^), tree shrew^50^ (73 mm^2^), and gray squirrel^35,38^ (82 mm^2^). Based on the tree shrew model (784 hidden units) we defined earlier, we set 900 units for gray squirrel, 25 units for the mouse and 64 units for the rat models.

## Supplementary Information

**Supplementary Fig.1.**
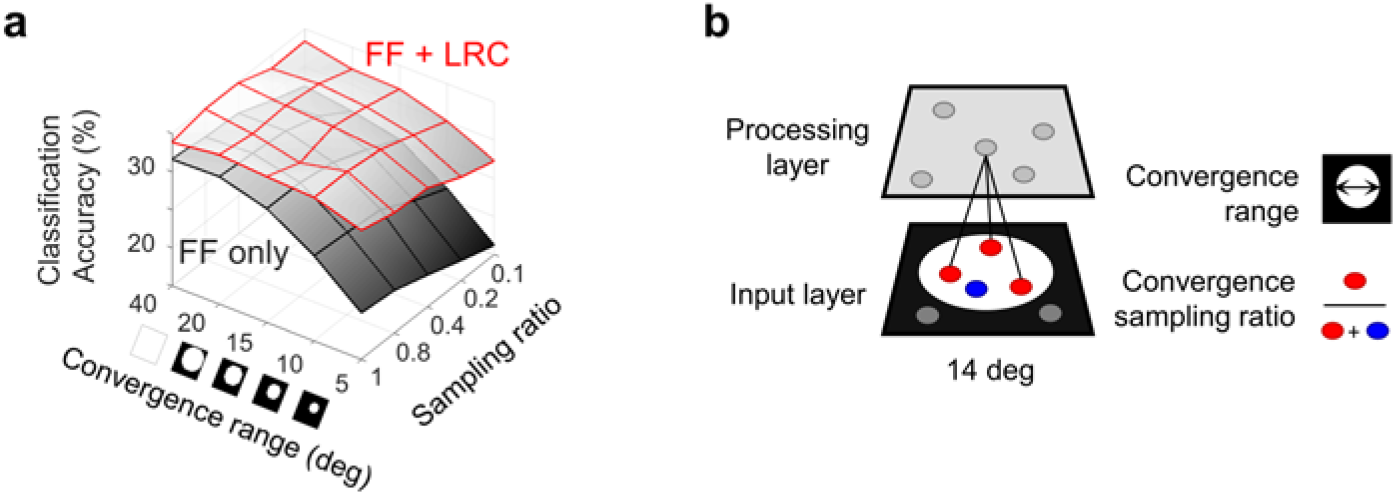
Effect of the LRCs on various feedforward connectivity. (a) Classification accuracy of the network for CIFAR-10 datasets in relation to the variation of structural parameters of the feedforward projections. The black surface represents performance of the feedforward network (FF only), and the red surface represents performance of the network with n=10,000 (1.2% of connections of fully connected networks in Fig 2) of LRCs added (FF + LRC). Performance is improved by LRCs, regardless of the condition of feedforward connectivity. (b) The convergence range of feedforward projections is defined as the diameter of sampling in the input layer. In the current model, 1 pixel of CIFAR-10 image was scaled to 0.5 degree in visual space. The sampling ratio is defined as the connection probability of neurons in the input layer within the convergence range.

**Supplementary Fig.2.**
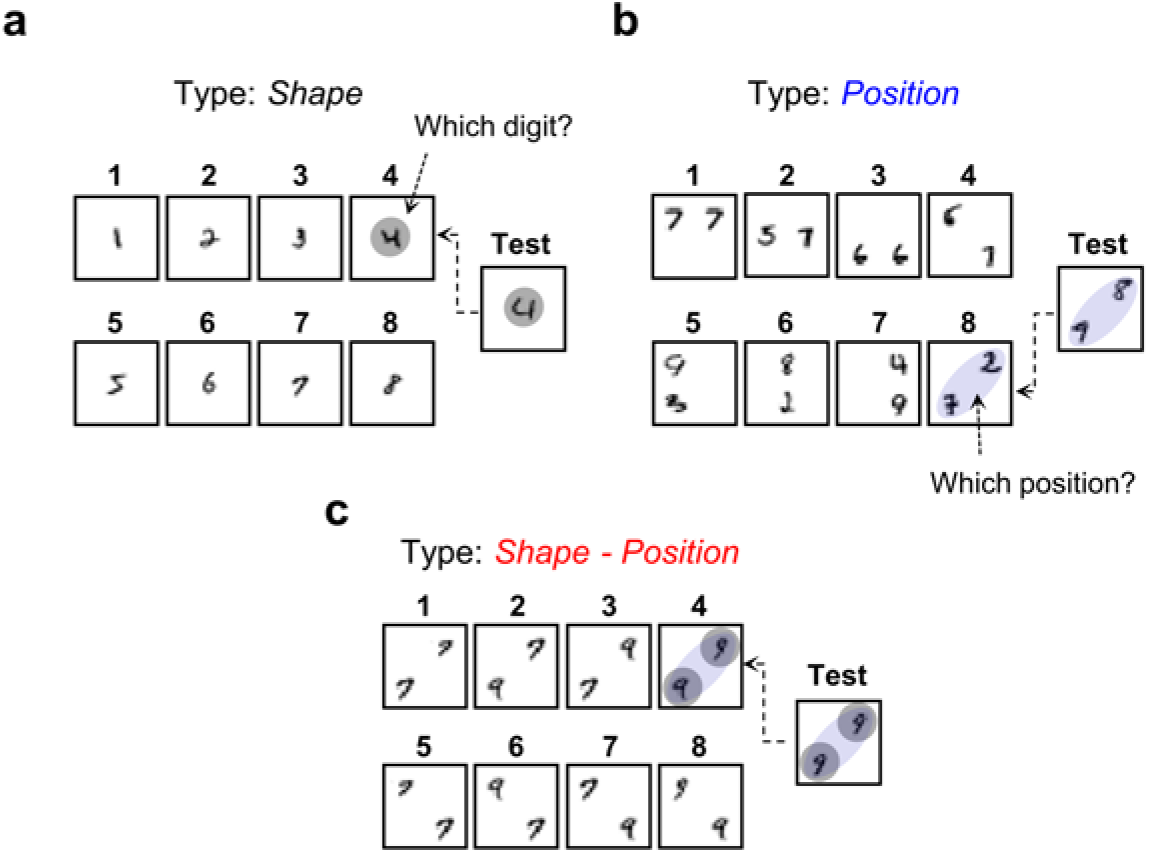
Modified MNIST datasets for the test of local and global feature classification. (a) The “shape” dataset was designed by arranging a hand-written digit of 8 × 8 pixels in the center of a 28 × 28-pixel image. The dataset consists of eight categories by value of the number in the center, chosen randomly from 1 to 8. (b) The “position” dataset was made using the following procedure. First, two digits of 8 × 8 pixels were randomly chosen. Second, these two digits were allocated in a 28 × 28 image, with one of the following position alignments: horizontal (top, middle, bottom), vertical (left, middle, right), or diagonal (45°, 135°). This dataset also consists of eight categories only depending on the positions of digits. (c) The “shape-position” dataset was made by the following procedure. First, a 28 × 28-pixel area was divided into four 14 × 14 areas. Second, two diagonally aligned areas were selected (either 45° or 135°). Third, one of two digits, either “7” or “9”, composed of 8 × 8 pixels was inserted into each selected area. This dataset consists of eight categories depending on both the shape and position of the digits.

**Supplementary Fig.3.**
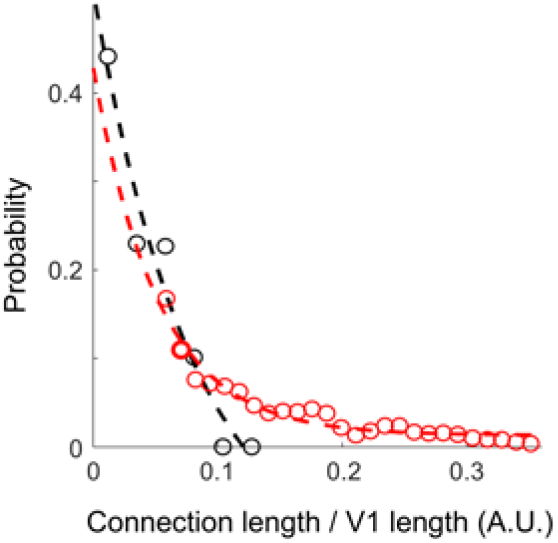
Length of lateral connections in tree shrews and mice. Distribution of the length of lateral connections in tree shrews (red circles; Bosking 1997) and mice V1 (black circles; Seeman 2018), normalized by the size of each V1.

**Supplementary Fig.4.**
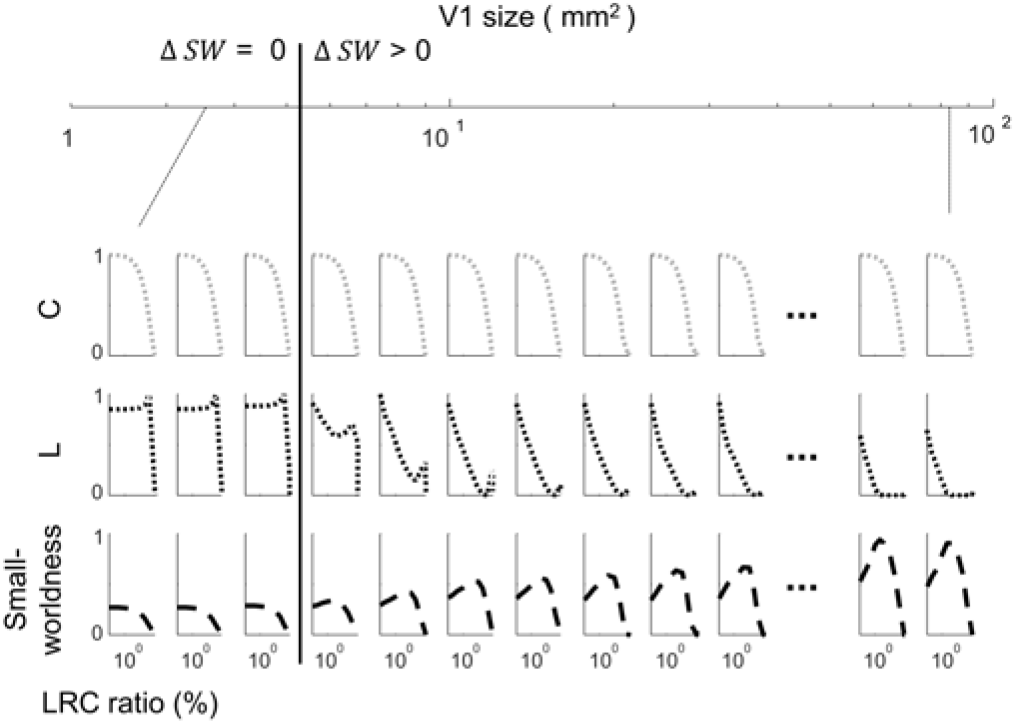
Changes in the network coefficients and small-worldness in relation to cortex size. Change of the cluster coefficient (C), average shortest path length (L), and small-worldness during the rewiring process for V1 of various sizes.

